## Supplementary Information for "Efficient Bayesian inference for mechanistic modelling with high-throughput data"

### Supplementary Information: Efficient Bayesian inference for mechanistic modeling with high-throughput data

#### S1 Data processing: removing data artifacts

##### S1.1 Presence of both wound edges and final condition

Owing to the large number of samples in the high-throughput experiment, there exists large variations in the wound locations recorded in Williams *et al.*'s data set [1]. While the algorithm described in the main text is well-placed to incorporate a variable wound location, measurements where the image of the initial condition fails to capture one of the wound edges are problematic for the implementation. When one (or both) wound edges are absent, the model will simulate cell trajectories as if no cells are present on that side of the wound, meaning that for the simulated final condition there will be no population of cells on that side of the wound. However, experimental data can potentially show outgrowth into the field of view, leading to a large discrepancy between observed and simulated data – even when the model would be able to capture the dynamics on one of the edges of the wound well. To mitigate this effect, we use the subset of the data in which both edges of the wound are present. We also screen so that only observations with both an initial and final condition are considered. From the total of 332 genetic knockdowns present in our data of Williams *et al.'s* high-throughput screen, applying both filters yields 118 candidates for investigation.

##### S1.2 Presence of extruded cells in the wound area

After having selected for the samples in which there is an initial and final condition as well as containing both wound edges, we use a wound mask to remove extruded cells in the wound area. During the experiment, some very isolated cells appear in the centre of the wound. These cell locations are recorded by the DeepScratch protocol [2], but the cells identified are dead and have been extruded by the tissue. Hence, we apply a mask to filter out the locations of extruded cells. For each sample, we compute a mask with the outline of the wound, and remove all cell locations that are on the inside of the wound.

### S2 ABC-SMC parameter sampling

#### S2.1 Minibatch-ABC

In this section we describe in detail the ABC algorithms used for inference in the main text. As noted in the main text, the idea behind ABC-SMC is to propagate a sample from the prior distribution, $\boldsymbol{\pi}(\theta)$, through a sequence of intermediate distributions towards the target distribution, $P(\boldsymbol{\theta} \mid \mathcal{D}_{\text{obs}})$. The intermediate distributions are typically a sequence of ABC approximations, $P_t(\boldsymbol{\theta} \mid \mathcal{D}_{\text{obs}})$ for $t \in \{1, \ldots, T\}$. We use a sequential importance sampling approach to ABC-SMC [3, 4, 5], also called population Monte Carlo. Each Monte Carlo sample $\{\boldsymbol{\theta}_n^{(t)}, w_n^{(t)}\}$ built at generation $t$ is used to construct an importance distribution by perturbing the Monte Carlo samples with a kernel $K(\bullet \mid t)$. This importance distribution is then used to generate the next generation's Monte Carlo sample using importance sampling, detailed in Algorithm 1. The final sample, $\{\boldsymbol{\theta}_n^{(T)}, w_n^{(T)}\}$ forms the output of the ABC-SMC algorithm. The complete ABC-SMC algorithm is summarised in Algorithm 2.

**Algorithm 1:** Minibatch-importance sampling

**Input:** Data, $\mathcal{D}_{\text{obs}} = \{y^i_{\text{obs}}\}_{i=1}^N$; number of parameters to sample, $N_{\text{gen}}$; number of parameters to accept, $N_{\text{acc}}$; batch size, $N_{\text{bs}}$; prior distribution, $\pi(\boldsymbol{\theta})$; importance distribution, $\hat{q}(\boldsymbol{\theta})$; model, $f(\bullet \mid \boldsymbol{\theta})$; distance function, $d$.

**Output:** $N_{\text{acc}}$ weighted samples $\{\boldsymbol{\theta}_n, w_n\}_{n=1}^{N_{\text{acc}}}$.

```
for n = 0, ..., N_gen do
    Increment n ← n + 1;
    Generate parameter vector θ_n according to given importance distribution q̂(θ) ;
    Set ε_n = 0;
    for i = 1, ..., N_bs do
        Draw data index ξ uniformly at random from {1, ..., N};
        Simulate y_sim ~ f(• | θ_n, y^ξ_obs(0));
        Update ε_n = ε_n + d(y_sim, y^ξ_obs);
    end
end
From {θ_n, w_n}_{n=1}^{N_gen}, select the N_acc parameters with lowest value of ε_n;
Set w_n = π(θ_n)/q̂(θ_n) for all accepted parameters θ_n.
```

**Algorithm 2:** Sequential Monte Carlo ABC

**Input:** Data, $\mathcal{D}_{\text{obs}} = \{y^i_{\text{obs}}\}_{i=1}^N$; number of parameters to generate, $N_{\text{gen}}$; number of parameters to accept, $N_{\text{acc}}$; batch size, $N_{\text{bs}}$; prior distribution, $\pi(\boldsymbol{\theta})$; importance distribution, $\hat{q}(\boldsymbol{\theta})$, proportional to $q(\boldsymbol{\theta})$; model, $f(\bullet \mid \boldsymbol{\theta})$; distance function, $d$; number of generations, $T$.

**Output:** Weighted samples $\{\theta_n^{(T)}, w_n^{(T)}\}_{n=1}^{N_T}$.

Set $q_1 = \pi$;
**for** $t = 1, \dots, T-1$ **do**
  Produce weighted samples $\{\theta_n^{(t)}, w_n^{(t)}\}_{n=1}^{N_t}$ using Algorithm 1 (ABC-IS) with importance distribution $\hat{q}_t$, number of parameters to generate $N_{\text{gen}}$, number of parameters to accept $N_{\text{acc}}$, batch size $N_{\text{bs}}$, model $f$ and distance $d$;
  Define the importance distribution, $\hat{q}_{t+1}$, proportional to:

$$q_{t+1}(\theta) = \begin{cases} \sum_{n=1}^{N_t} w_n^{(t)} K_t(\boldsymbol{\theta} \mid \boldsymbol{\theta}_n^{(t)}) / \sum_{m=1}^{N_t} w_m^{(t)} & \text{if } \pi(\boldsymbol{\theta}) > 0 \\ 0 & \text{else;} \end{cases}$$

**end**

#### S2.2 Choice of knockdown kernel

In this work, we use the multivariate normal kernel proposed by Filippi *et al.* [6], which is designed to approximate the target density well, while also maximizing the acceptance rate at each step [6]. In effect, this kernel perturbs each sampled particle according to a multivariate distribution with covariance matrix $\Sigma^{(t)}$, which depends on the covariance of the previous population. Filippi *et al.* [6] prove that the optimal choice of kernel is then approximatly

$$\Sigma^{(t)} \approx \sum_{k=1}^{N_{\text{acc}}} \sum_{\ell=1}^{N_{\text{acc}}} w_k^{(t-1)} w_\ell^{(t-1)} (\boldsymbol{\theta}_k - \boldsymbol{\theta}_\ell)(\boldsymbol{\theta}_k - \boldsymbol{\theta}_\ell)^T. \tag{S1}$$

This allows us to define the proposal mechanism $K(\bullet|\boldsymbol{\theta}_n^{(t)})$ defined in Algorithm 2, as

$$K(\bullet|\boldsymbol{\theta}_n^{(t)}) \sim \mathcal{N}(\boldsymbol{\theta}_n^{(t)}, \Sigma^{(t+1)}), \tag{S2}$$

where $\Sigma^{(t+1)}$ is defined in Equation (S1).

#### S2.3 Model calibration: number of ABC-SMC generations

For each ABC-SMC implementation of this specific model, we must define the number of generations, $T$. This is an additional hyperparameter whose choice requires further justification.

Choosing a higher number of generations will be computationally more costly, while choosing a lower number of generations risks not approximating the true posterior accurately. To balance between these two requirements, we perform test runs of minibatch ABC-SMC with our chosen batch size of $N_{\text{bs}} = 10$ with a varying number of generations to quantify the gain of information after each subsequent generation. One way to measure this gain is the Kullback-Leibler (KL) divergence [7] between each generation. The KL divergence for two probability distributions, $p$ and $q$, defined on a common space $\Omega$, is defined as

$$\mathrm{KLDiv}(p\|q) = \int_{\Omega} p(x) \log\left(\frac{p(x)}{q(x)}\right) \mathrm{d}x. \tag{S3}$$

The KL divergence as a measure of change of the distribution between one generation and the next has been used previously by Filippi *et al.* [6] to derive kernels that optimally increase the difference between two generations. Note that we also use the resulting kernel proposed by Filippi *et al.* [6]. In this work, we similarly consider how the quantity

$$d_t = \mathrm{KLDiv}(p_{t+1}\|p_t), \tag{S4}$$

evolves over time, where $p_t$ is the ABC-SMC posterior after $t$ generations. We plot $d_t$ for the three genetic knockdowns for which most data is available (Mock, CDC42, and CDH5 knockouts) in Figure S1 to find that including additional generations after $T = 4$ generations have virtually no impact on the posteriors, hence we choose this cut-off for the number of generations.

#### S3 Justification of model reduction

We explore the impact of adding a density-dependent term to contribute to the rate of cell death, analogous to those used to model cell movement and proliferation. This gives rise to two competing models. In the simpler model, death occurs at a constant rate, $d$, independent of local crowding. In the more complex model, the density-dependent death rate for agent $n$, $D_n$, is defined by

$$D_n = \max\left\{0, d - \gamma_d \sum_{i=1, i\neq n}^{N(t)} \exp\left(\frac{-\|\boldsymbol{x}n - \boldsymbol{x}_i\|^2}{2\sigma^2}\right)\right\}. \tag{S5}$$

To quantify whether the data contain evidence in favour of one model as opposed the other, we compute the Bayes factor between the simple (density-independent) model, $M_{\text{simple}}$, and the complex (density-dependent) model, $M_{\text{complex}}$. The Bayes factor is then defined as

$$\mathcal{B} = \frac{\mathbb{P}(\mathcal{D}_{\text{obs}}|M_{\text{simple}})}{\mathbb{P}(\mathcal{D}_{\text{obs}}|M_{\text{complex}})}, \tag{S6}$$

and the value of $\mathcal{B}$ quantifies the strength of the evidence in favour of $M_{\text{simple}}$ [5]. A value of $\mathcal{B}$ between 1 and 3 represents very weak evidence in favour of $M_{\text{simple}}$, a value between 3 and

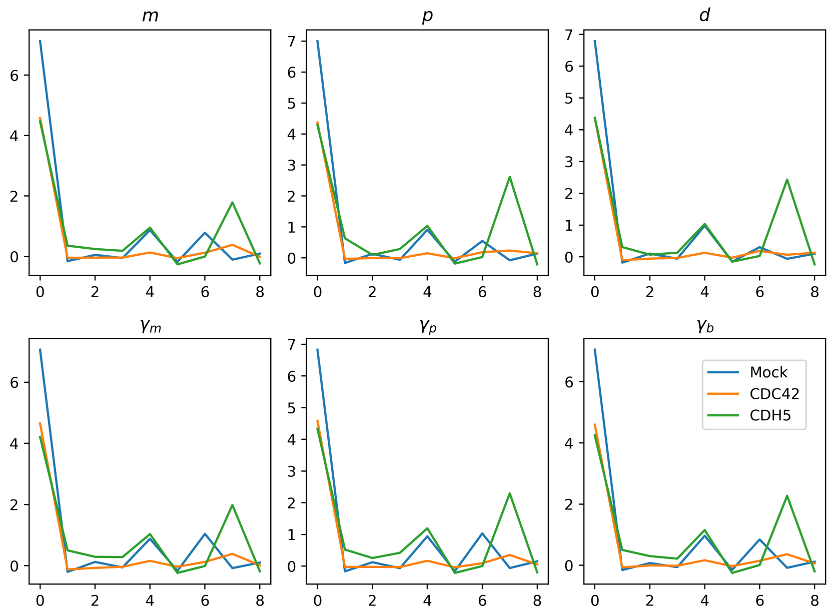


Figure S1: KL divergence between the posterior distributions at one generation and the next, plotted for a total of 10 generations. The KL divergence has decreased significantly after just two generations as compared to the prior and subsequent generations do not diminish the KL divergence substantially.

20 represents positive evidence, between 20 and 150 strong evidence and above 150 very strong evidence [8]. From the ABC-SMC posterior distributions the Bayes factor for each model can be computed straightforwardly [9]. For a given set of population weights $\{w_i^t\}_{i=1,\ldots,n,t=1,\ldots,T}$, where $n$ is the number of samples in each generation, and $T$ is the number of generations,

$$\mathbb{P}(\mathcal{D}_{\text{obs}}|M) = \prod_{t=1}^{T} \frac{1}{n} \sum_{i=1}^{N} w_t^i. \tag{S7}$$

For the Mock, CDC42 and CDH5 knockout cases, we compute the Bayes factor, $\mathcal{B}$, using the simple and complex models. This gives $\mathcal{B} = 10.7, 4.69, 4.34$ for Mock, CDC42 and CDH5 knockouts, respectively, meaning that the data provides positive evidence in favour of the simpler model.

#### S4 Reliable inference using small batch sizes

The introduction of minibatches in ABC-SMC comes with the task of selecting the batch size as an algorithm hyperparameter. A good choice for the batch size, $N_{\text{bs}}$, needs to take into account the balance between computational complexity on the one hand and a reliable estimate of the discrepancy between model and data on the other hand. In general, the capacity of any ABC implementation to infer parameter values from data will depend on the specific computational model and the data available (for instance, through its spatial and temporal resolution, or its signal to noise ratio). To investigate the performance of minibatch-ABC-SMC in identifying parameters from data in our specific context, we define two different parameter regimes and assess the performance of minibatch-ABC-SMC in identifying the parameter values on a synthetic dataset created using the parameter values in the two regimes. We let parameter regime I correspond to a high movement rate with a positive impact of density on movement and a low proliferation rate, and choose $m = 1.5h^{-1}$, $p - d = 0.01h^{-1}$, $\gamma_m = -1h^{-1}$, $\gamma_p = 0.01h^{-1}$ and $\gamma_b = 20\mu m$[1]. We let parameter regime II correspond to a low movement rate with contact inhibition for movement and a high proliferation rate, and choose $m = 0.5h^{-1}$, $p - d = 0.025h^{-1}$, $\gamma_m = 1.5h^{-1}$, $\gamma_p = 0.01h^{-1}$ and $\gamma_b = 20\mu m$[2].

##### S4.1 Variability of the posterior mean

For each parameter regime, an *in silico* data set is created by simulating one model output at $t = 24h$ for every initial condition corresponding to Mock. That is, each of the two *in silico* datasets contains 117 data points, each corresponding to one data point in the *in vitro* dataset from the high-throughput experiment. Then, we perform minibatch-ABC-SMC with each of the two synthetic data sets with varying batch size, such that $N_{\text{bs}} = 1, 5, 10, 50, 117$. For each parameter regime and batch size, we repeat the inference process five times to quantify the variability of the estimated posterior distributions. Figure S3 shows boxplots for the resulting posterior means for each of the parameter regimes as the batch size is increased. It is clear from Figure S3 that the performance of the algorithm in identifying the true parameter value (horizontal lines in the figure) is affected by variations in the batch size, $N_{\text{bs}}$. However, there is a critical value – in Figure S3 this is typically around $N_{\text{bs}} = 10$ – after which increasing the batch size, $N_{\text{bs}}$, does not reduce the variability of the estimated posterior means. Similarly, after this critical value, the capacity of minibatch-ABC-SMC to identify the true posterior mean does not improve, even when the batch size is increased to the full data size. At any value of the batch size, $N_{\text{bs}}$, the minibatch-ABC-SMC implementation is capable of correctly discerning structural differences between the two parameter regimes. For instance, for the data generated using parameter regime I, the estimated intrinsic motility rate, $m$, is consistently estimated as much higher than that of the data from parameter regime II, as is consistent with the definition of the parameter regimes. Conversely, the net intrinsic proliferation rate, $p - d$, is estimated as much higher in parameter regime II than in parameter regime I. For the density-dependent interaction term, $\gamma_m$, the minibatch ABC-SMC implementation can discern at any batch size

[1] To sample $p - d = 0.01h^{-1}$, we sample $p \in (0, 0.05)$ and then choose $d = 0.01 - p$.

[2] $p$ and $d$ are sampled analogously as in parameter regime I.

that the effect of crowding on motility is positive in regime I, and negative in regime II.

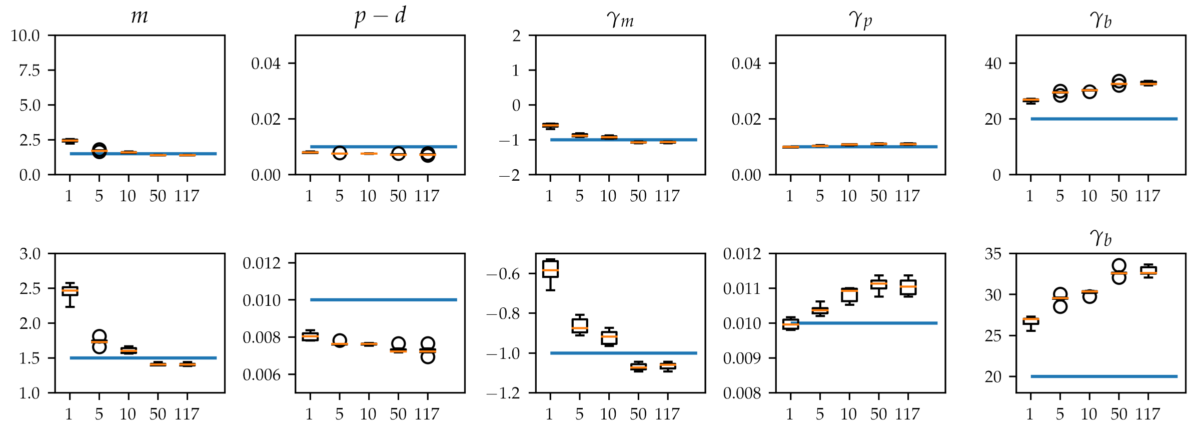


Figure S2: Boxplots for the means of the posterior distributions obtained by performing minibatch ABC-SMC on parameter regime I when the batch size is varied. The blue horizontal line indicates the true parameter value. Top row: boxplots over the full prior range. Bottom row: zoomed range to more accurately display the discrepancy between estimates and the true parameter value.

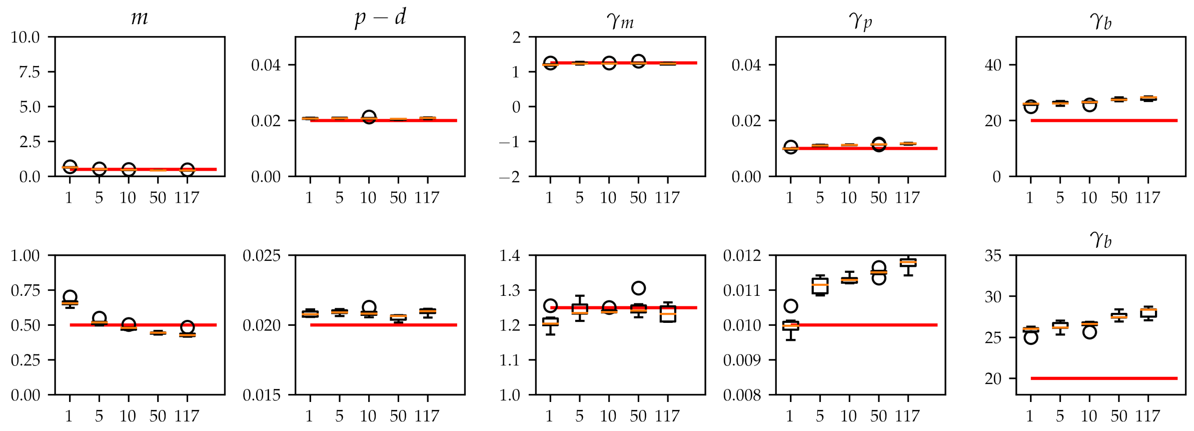


Figure S3: Boxplots for the means of the posterior distributions obtained by performing minibatch-ABC-SMC on parameter regime II when the batch size is varied. The red horizontal line indicates the true parameter value. Top row: boxplots over the full prior range. Bottom row: zoomed range to more accurately display the discrepancy between estimates and the true parameter value.

#### S4.2 Variability of the posterior variance

To further assess the similarity of the learned posterior distributions when the batch size, $N_{\text{bs}}$, is varied, we choose one of the posterior distributions for each of the experiments using each of $N_{\text{bs}} = 1, 5, 10$ and show the distributions in Figure S4. The behaviour of the posterior distributions in Figure S4 is in agreement with the idea that the qualitative behaviours of the model can be estimated well even at small batch size, as even the smallest batch sizes provide posterior distributions with posterior means close to the true value used to generate the synthetic data. It is clear from Figure S4 that increasing the batch size, $N_{\text{bs}}$, helps to decrease the spread of the posterior distribution.

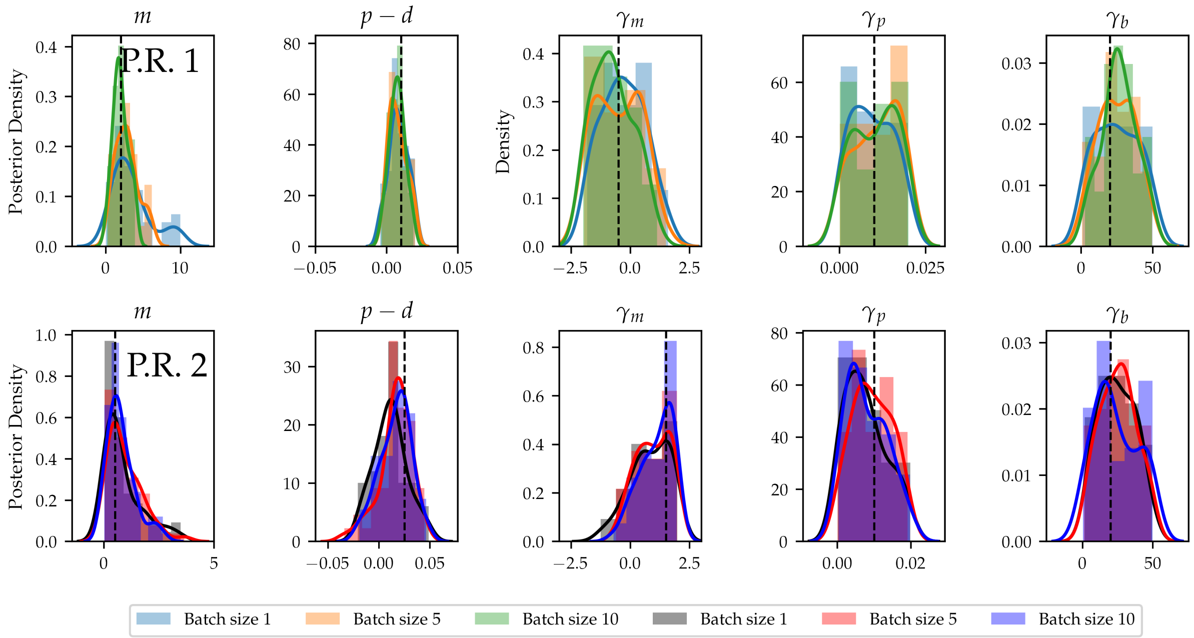


Figure S4: Posterior distributions obtained using the two synthetic data sets corresponding to parameter regimes I and II. In all plots, the dashed black line indicates the true parameter value. Top row: posterior distributions for the data set generated using parameter regime I. Bottom row: posterior distributions for the data set generated using parameter regime II.

Figure S4 suggests that increasing the batch size, $N_{\text{bs}}$, reduces the variance of the posterior distribution. To quantify this effect, we analyse the standard deviation of the learned posterior distributions at each different batch size. The standard deviation of the posterior density serves as a useful measure to quantify the uncertainty associated with the posterior distribution. Figure S6 reveals that for three key parameters in the model, the standard deviation decreases with an increasing batch size. Strikingly, the standard deviation drops rapidly when the batch size is increased initially, but then stays relatively constant across larger batch sizes. In both parameter regimes, this happens for three of the parameters around $N_{\text{bs}} = 10$, while for the remaining parameters the posterior standard deviation remains roughly constant when the batch size is increased further. Taken together, the analysis of posterior means and posterior standard deviation suggests that a minibatch implementation of ABC-SMC with small batch sizes can have a similar performance to conventional ABC-SMC (that uses very large batch sizes), while the number of simulations needed is much lower (in this case, 10 simulations per sampled parameter vector instead of 117).

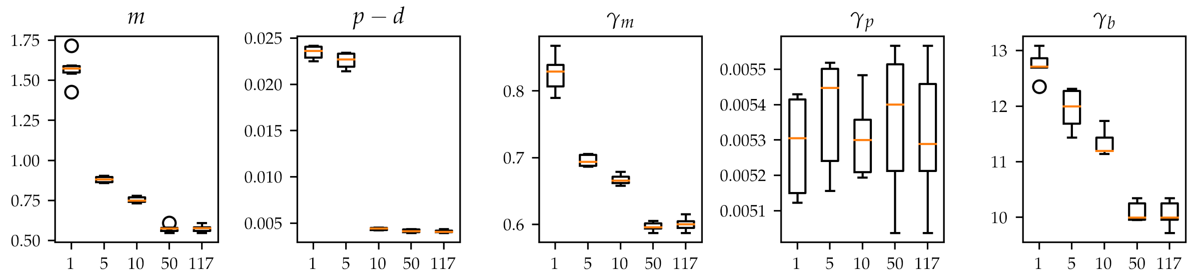


Figure S5: Boxplots for the standard deviations of the posterior distributions obtained by performing minibatch-ABC-SMC on parameter regime I when the batch size is varied.

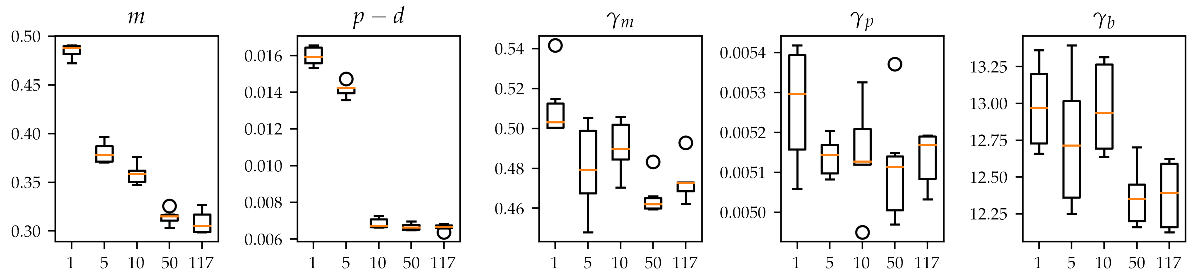


Figure S6: Boxplots for the standard deviations of the posterior distributions obtained by performing minibatch-ABC-SMC on parameter regime II when the batch size is varied.

#### S4.3 Performance of different batch sizes on experimental data

Taken together, our analysis shows that on synthetic data, a minibatch-ABC-SMC implementation has comparable performance to a traditional ABC-SMC implementation in retrieving a known model parameter value and is capable of inferring meaningful mechanistic insights using the parametrised model. To explore the effect that a small batch size has on the qualitative properties of the posterior distribution when applied to real data, we perform inference on the mock data set, where we vary the batch size from 1 to 117. In Figure S7, we compare the posterior distributions. Figure S7 shows that the learned posterior distributions are qualitatively very similar, meaning that the intrinsic variability of the data is captured well by batch sizes as small as $N_{\text{bs}} = 10$. Taken together with the uncertainty quantification carried out for the two parameter regimes that are used to create the synthetic data, we propose to use a batch size $N_{\text{bs}} = 10$ for the analysis of images for which a large number of observations are recorded in

Williams *et al.*'s mRNAi screen [1].

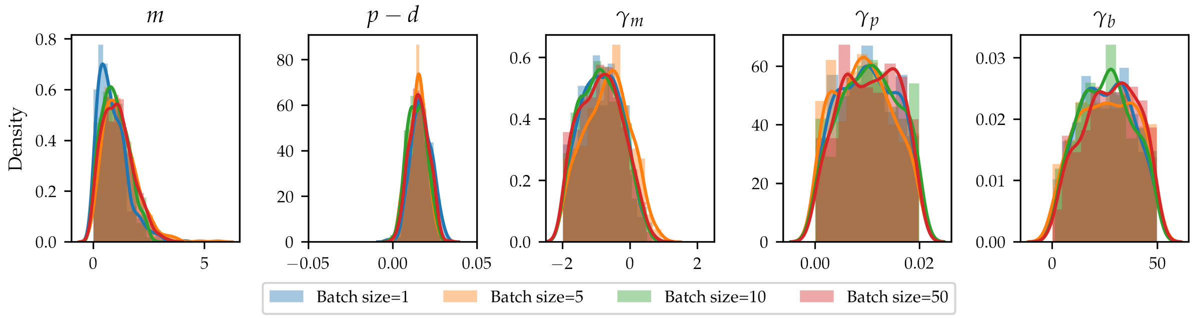


Figure S7: Posterior densities of model parameters for the mock data set. When the batch size, $N_{\text{bs}}$, is increased, the posteriors remain qualitatively very similar and consistently identify the same maximum a posteriori parameter value.

#### S4.4 Capacity of inference with very small batch sizes on experimental data

Our previous experiments suggest that for this specific model and data, the choice of $N_{\text{bs}} = 10$ offers a good balance between computational complexity, robust inference of the posterior mean and low variance of the posterior distribution. Decreasing the batch size to $N_{\text{bs}} = 2$ still offers reliable identification of the posterior mean, albeit at the cost of a higher variance in the posterior distribution. Since many of the knockdowns in the high-throughput data set contain only two datapoints, we wish to understand if the posterior distributions obtained by applying minibatch-ABC-SMC when only two datapoints are available can still lead to the identification of mechanistically relevant differences between the knockdowns. To do this, we select 40 datapoints from each of the Mock and CDH5 knockdown experiments at random and without replacement, and divide them into 20 pairs of two observations for each of the datasets. We perform minibatch-ABC-SMC on each of the resulting *miniature data sets* and record their posterior distributions. Figure S8 shows boxplots for the posterior distributions for each of the Mock and CDH5 knockdown experiments.

From Figure S8 it can be seen that while the spread of the parameter values is indeed larger for each of the experimental conditions than when the batch size is larger, the boxplots of the posterior means suggest that meaningful differences between the two different genetic knockdowns can be identified. This suggests that even a very small number of observations can be used to identify the posterior distributions of model parameters given data in the high-throughput experiment.

To compare the performance of minibatch ABC-SMC with naive subsampling from the data, we fix $N_{\text{bs}} = 10$ as in the main text and investigate the quality of the posterior distributions ob-

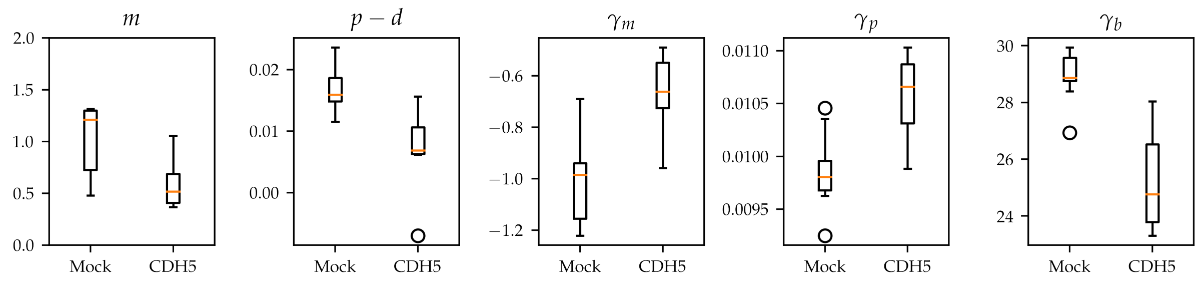


Figure S8: Boxplots showing the means of the posterior distributions found by performing minibatch-ABC-SMC on *miniature data sets* created by taking subsets of the data. The boxplots show that there are meaningful separations of the posterior means between the mock and CDH5 knockdowns, even when each *miniature data set* consists of only two datapoints.

tained by performing ABC-SMC on subsamples of size $N_{\text{bs}}$ of the data. From the Mock dataset, we select 100 datapoints at random and without replacement, and divide them into 10 subsets of $N_{\text{bs}} = 10$ datapoints. As in Supplementary Information Section 4.4, we perform minibatch-ABC-SMC on each of the resulting *miniature data sets* and record their posterior distributions. We independently run 10 instances of of minibatch ABC-SMC, again using $N_{\text{bs}} = 10$, and record the posterior distributions. Figure S9 shows boxplots of the means of the posterior distributions found by performing ABC-SMC on subsets of the data as well as using minibatch ABC-SMC. The boxplots show that the subsample approach yields more variable posterior means than minibatching.

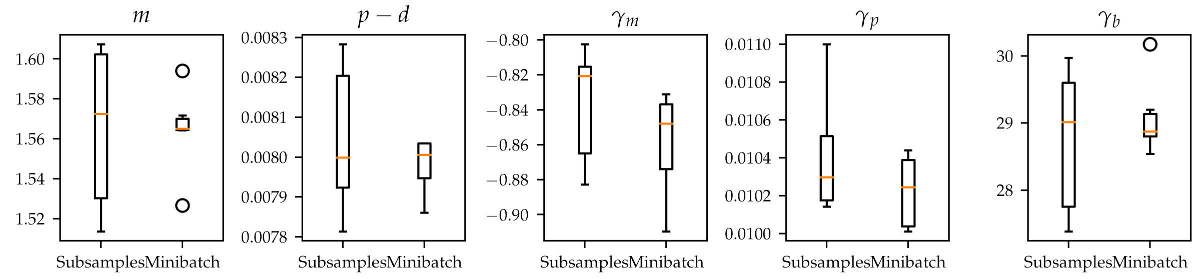


Figure S9: Boxplots of the means of the posterior distributions found by performing ABC-SMC on subsets of the data as well as using minibatch-ABC-SMC. Subsampling yields more variable posterior means than minibatching.

Figure S10 shows boxplots of the variance of the posterior distributions generated by performing ABC-SMC on subsets of the data as well as using minibatch-ABC-SMC. The boxplots show that the subsample approach yields on average larger posterior variance than minibatching, indicating that the posterior obtained using subsampling contains more uncertainty than that obtained using minibatch ABC-SMC.

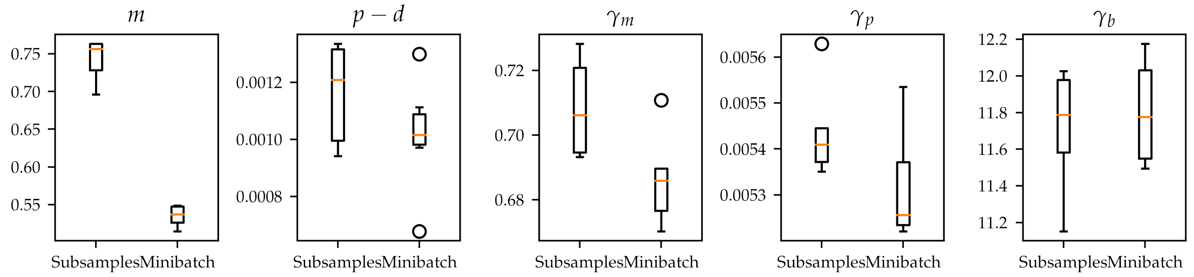


Figure S10: Boxplots of the variances of the posterior distributions found by performing ABC-SMC on subsets of the data as well as using minibatch-ABC-SMC. Subsampling yields larger posterior variances than minibatching.

#### S5 Robustness of model predictions to cell size esimates

An important model parameter in the IBM is the cell size, which is used to define the exponential kernel that governs cell-cell interactions. Within each genetic knockdown, the cell area is estimated using the protocol of Javer *et al.* [2] for each each individual observed cell. In Figure S11 we show the distribution of cell areas across all different genetic knockdowns (for a total of approximately $5 \cdot 10^5$ cells. We see that the area distribution is sharply peaked, with some very large outliers corresponding to errors in the Voronoi tesselation of Javer *et al.* [2]. For this reason, we consider only cells whose areas are measured as smaller than 500 square pixels.

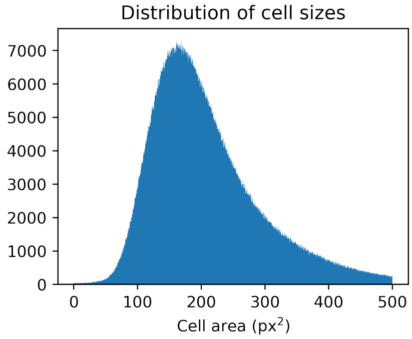


Figure S11: Distribution of cell sizes in the dataset.

For each knockdown, we estimate the cell area parameter for simulations as the median of the cell area measurements for that knockdown. To address the robustness of our model predictions to errors in cell area measurement, we carry out uncertainty quantification analysis by varying the cell area input parameter for both mock and CDH5 knockdowns. For each of mock and

CDH5 knockdown, we simulate from the model using a range of cell area measurements defined by variations of up to 50% of the cell area median. This corresponds roughly to simulating within five standard deviations of the input parameter. For each value in the cell area range, we simulate the model ten times. In Figure S12, we plot how much the resulting posterior mean changes relative to the posterior mean obtained when using the measured cell area parameter, *i.e.* for $\mu$ the posterior mean obtained by using the cell area input parameter used in the main text, and $\mu^i_{A_j}$ the $i$-th posterior mean obtained by using cell area $A_j$, we estimate the relative error for cell size $A_j$, $\epsilon_{\text{rel}}$, as

$$\epsilon_{\text{rel}} = \frac{\frac{1}{10}\sum_{i=1}^{10} |\mu^i_{A_j} - \mu|}{|\mu|}, \tag{S8}$$

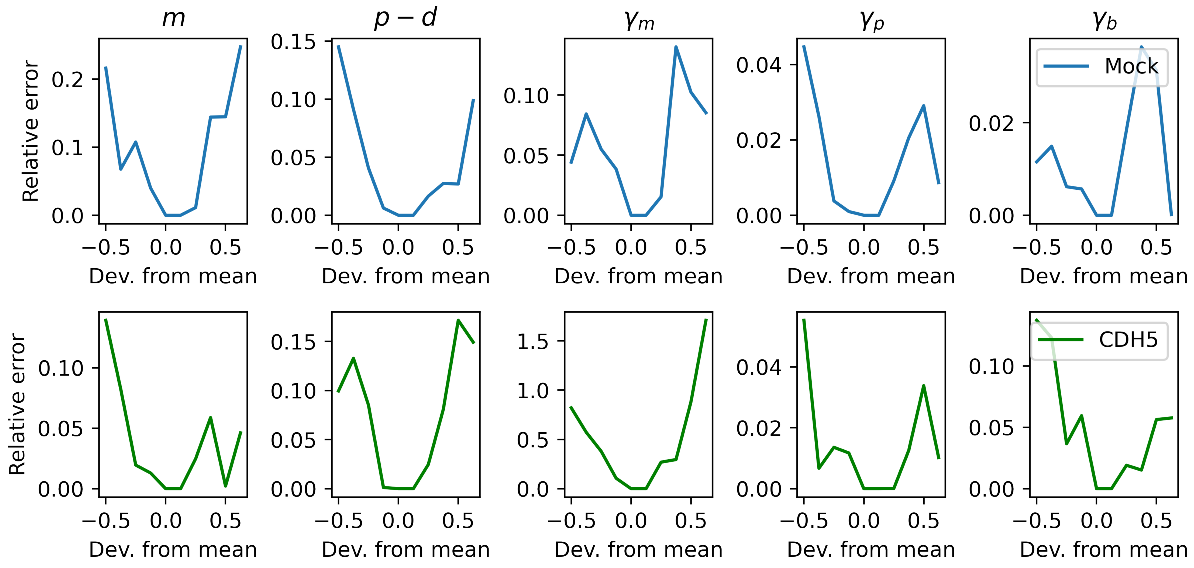


Figure S12: Relative error plotted against deviations of the cell area as input parameters for minibatch-ABC-SMC. Deviations within roughly three standard deviations from the median result in the relative error in the posterior mean below 10%.

We find that for cell areas within 25% of the median cell area, which roughly corresponds to cell areas within three standard deviations from the median, the relative error in the posterior mean is below 10% for both the mock and CDH5 knockdown, from which we conclude that our predictions are robust to small errors in the estimation of cell areas.

#### S6 Changing the number of clusters for $K$-means clustering

In the main text, $K$-means clustering is performed on the posterior means of the different knockdowns for $K = 3$. In this section, we show the sum of squared distances when $K$ is varied in the range $1, \ldots, 10$ to find that marginal improvement in sum of squares rapidly diminishes above $K = 3$, suggesting that $K = 3$ is a good choice for the number of clusters in the analysis.

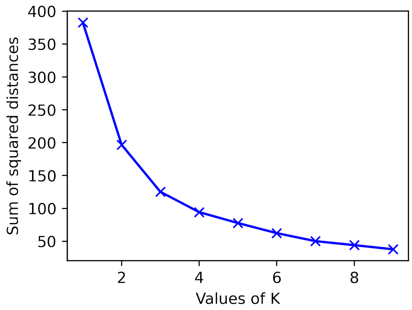


Figure S13: Sum of squared distances for $K$-means clustering as $K$ is increased. The marginal improvement of the sum of squared distances markedly decreases after $K = 3$, suggesting that this is the optimal choice for the number of clusters.

#### S7 Confidence intervals for model parameters

In this section, we report the confidence intervals computed for the experiments in main text Section 3.2. For each experiment, we compute the 99% confidence interval, which for a sample of size $n$ of a random variable $x$, with sample mean $\bar{x}$ and sample standard variance $\sigma$ under the normality assumption, is given by

$$\mathrm{CI} = \left( \bar{x} - z\frac{\sigma}{\sqrt{n}}, \bar{x} + z\frac{\sigma}{\sqrt{n}} \right), \tag{S9}$$

where $z$ equals the $z$-score, which for a 95% confidence interval is $z = 1.96$.

##### S7.1 Confidence intervals for posterior means of Mock, CDC42, CDH5 knockdowns

| Experiment | Posterior mean | Confidence interval |
|---|---|---|
| Mock, $m$ | 0.7194 | (0.6789,0.7599) |
| Mock, $p-d$ | 0.01493 | (0.01448, 0.01538) |
| Mock, $\gamma_m$ | -1.06206 | (-1.1092, -1.01493) |
| Mock, $\gamma_p$ | 0.01014 | (0.009652, 0.01062) |
| Mock, $\gamma_b$ | 29.1372 | (28.08166, 30.1927) |
| CDC42, $m$ | 0.3278 | (0.3054, 0.3501) |
| CDC42, $p-d$ | 0.01089 | (0.01030, 0.01147) |
| CDC42, $\gamma_m$ | -0.2915 | (-0.3121, -0.2709) |
| CDC42, $\gamma_p$ | 0.009133 | (0.008648, 0.009618) |
| CDC42, $\gamma_b$ | 25.4509 | (24.2694, 26.6324) |
| CDH5, $m$ | 0.43168 | (0.40364, 0.4597) |
| CDH5, $p-d$ | 0.005825 | (0.005353, 0.0062966) |
| CDH5, $\gamma_m$ | -0.6170 | (-0.6524, -0.5816) |
| CDH5, $\gamma_p$ | 0.01069 | (0.01022, 0.01116) |
| CDH5, $\gamma_b$ | 27.02407 | (25.8691, 28.1790) |

Table S1: Posterior means and confidence intervals of posterior means for each of the experiments.

##### S7.2 Confidence intervals for posterior means of different initial wound sizes

| Experiment | Posterior mean | Confidence interval |
|---|---|---|
| Mock, small wound, $m$ | 1.4164 | (1.3403, 1.4926) |
| Mock, middle wound, $m$ | 0.6873 | (0.6506, 0.7242) |
| Mock, large wound, $m$ | 0.8639 | (0.8168, 0.9111) |
| Mock, small wound, $p-d$ | 0.01869 | (0.01825, 0.01913) |
| Mock, middle wound, $p-d$ | 0.01479 | (0.01436, 0.01522) |
| Mock, large wound, $p-d$ | 0.01462 | (0.01415, 0.01509) |
| Mock, small wound, $\gamma_m$ | -1.1068 | (-1.1620, -1.0513) |
| Mock, middle wound, $\gamma_m$ | -1.1668 | (-1.2102, -1.1234) |
| Mock, large wound, $\gamma_m$ | -1.0026 | (-1.05234, -0.95288) |
| Mock, small wound, $\gamma_p$ | 0.010966 | (0.01048, 0.01145) |
| Mock, middle wound, $\gamma_p$ | 0.01136 | (0.01089, 0.011835) |
| Mock, large wound, $\gamma_p$ | 0.009991 | (0.009506, 0.01047) |
| Mock, small wound, $\gamma_b$ | 30.9606 | (29.9807, 31.9406) |
| Mock, middle wound, $\gamma_b$ | 30.8423 | (29.8193, 31.8654) |
| Mock, large wound, $\gamma_b$ | 29.336 | (28.2649, 30.4070) |

Table S2: Posterior means and confidence intervals of posterior means of different initial wound sizes for Mock.

| Experiment | Posterior mean | Confidence interval |
|---|---|---|
| CDH5, small wound, $m$ | 0.4034 | (0.3738, 0.4330) |
| CDH5, large wound, $m$ | 0.3584 | (0.3352, 0.3815) |
| CDH5, small wound, $p - d$ | 0.006947 | (0.0064504, 0.007443) |
| CDH5, large wound, $p - d$ | 0.0068595 | (0.006376, 0.0073425) |
| CDH5, small wound, $\gamma_m$ | -0.7420 | (-0.7818, -0.7022) |
| CDH5, large wound, $\gamma_m$ | -0.8277 | (-0.8653, -0.7901) |
| CDH5, small wound, $\gamma_p$ | 0.01090 | (0.01042, 0.01139) |
| CDH5, large wound, $\gamma_p$ | 0.01088 | (0.01040, 0.01136) |
| CDH5, small wound, $\gamma_b$ | 27.366 | (26.2392, 28.4930) |
| CDH5, large wound, $\gamma_b$ | 29.1108 | (27.9956, 30.2259) |

Table S3: Posterior means and confidence intervals of posterior means of different initial wound sizes for CDH5 knockdown.

[2] Avelino Javer, Jens Rittscher, and Heba Sailem. Deepscratch: Single-cell based topological metrics of scratch wound assay. *Computational and Structural Biotechnology Journal*, 18, 08 2020.
